# Distilling the neurophenomenological signatures of pure awareness during Transcendental Meditation

**DOI:** 10.1101/2025.09.25.678494

**Authors:** Alejandro Chandia-Jorquera, Sean D. van Mil, Mar Estarellas, Marina Dauphin, Claudia Pascovich, Andres Canales-Johnson

## Abstract

Pure awareness (PA) has been proposed as a form of minimal phenomenal experience, but its neurophenomenological signatures remain poorly characterized. Transcendental Meditation (TM) offers a particularly tractable empirical model of PA because its procedure is standardized, its induction is effortless, and it reliably elicits reports of awareness with minimal content. We combined electroencephalography (EEG) with Temporal Experience Tracing in 33 experienced TM practitioners and their matched controls (performing mental counting). TM practitioners reported significantly greater intensity and temporal variability of PA, independent of years of meditation practice. We then used multivariate classification of theoretically motivated EEG markers spanning temporal entropy, aperiodic activity, complexity, and linear and nonlinear functional connectivity. We observed a double dissociation. When TM was contrasted with counting, temporal entropy and aperiodic dynamics were the strongest discriminators, whereas phase-coherence functional connectivity contributed least. Conversely, when TM was contrasted with its own baseline, low-frequency functional connectivity dominated, whereas temporal entropy contributed minimally. Complementary topographical analyses indicated that these differences were not reducible to a few localized univariate effects, but were better understood as distributed multivariate neural patterns. Finally, TM showed little evidence of carryover into subsequent rest, whereas counting induced more residual change. Together, these findings provide a systematic electrophysiological characterization of PA and support neurophenomenology as a tractable framework for studying minimal phenomenal experience.

## INTRODUCTION

Pure awareness (PA) refers to a phenomenological experience described in many contemplative traditions (Forman, 1986; Potash et al., 2025b; Travis and Pearson, 2000; Vieten et al., 2018). It designates a paradigmatic case of minimal phenomenal experience (MPE): an episode of wakeful experiencing stripped of customary structural features of consciousness (Metzinger, 2020). Phenomenologically, PA is marked by the absence of minimal phenomenal selfhood (i.e., no identification with a body or ego), the lack of an explicit temporal register, and the absence of a spatial frame of reference, while retaining a distinctive sense of wakefulness or alert presence (Forman, 1986; Metzinger, 2020; Travis, 2014; Travis and Pearson, 2000; Vieten et al., 2018).

Within the vast repertoire of meditation practices, Transcendental Meditation (TM) offers a tractable empirical model of PA as its procedure is standardized, its induction is effortless and repeatable, and its reported phenomenology closely matches the defining features of minimal phenomenal experience (Travis, 2014; Travis and Pearson, 2000). TM is a mantra-based practice, usually performed seated with eyes closed for 20 minutes twice a day, taught by certified teachers affiliated with the organization established by Maharishi Mahesh Yogi. During practice, TM practitioners begin by repeating a specific, nonsemantic sound (i.e., a ‘mantra’) that serves as a psychophysiological ‘vehicle’ for progressively attenuating discursive mentation. Although the mantra is not actively suppressed, it spontaneously recedes from awareness as attention settles. Practitioners frequently describe a shift from thought-mediated cognition to a state of pure awareness: awake but content-free, lacking an intentional object, devoid of deliberate mental activity, and minimally engaged with sensory input (Travis, 2014; Travis and Pearson, 2000). Crucially, TM is explicitly not framed as focused attention or open monitoring; instead, the mantra is used in a light, non-effortful manner, intended to allow attention to settle, a process described as ‘automatic self-transcendence’ (Travis et al., 2010). Thus, while the technique is initiated through the use of a mantra, its aim is the transcendence of the mantra itself, culminating in a putative mode of consciousness in which awareness is present with minimal representational content.

Despite the increasing adoption of neurophenomenological approaches in neuroscience of consciousness (Chowdhury et al., 2025; Ganesan et al., 2024; Jachs, 2021; Lewis-Healey et al., 2024b; Lutz, 2002; Potash et al., 2025b; Timmermann et al., 2023), a comprehensive understanding of how minimal phenomenal experiences, such as PA, relate to neural signatures remains lacking. This may be due to (i) the use of Likert scales to quantify the overall intensity of experiential dimensions, thus ignoring the temporal dynamics of phenomenology, and (ii) the lack of studies performing a large-scale screening of theoretically motivated neural markers that capture the complexity and nonlinear nature of brain dynamics subserving perception and consciousness (Canales-Johnson et al., 2020a, 2023, 2021b; Chidichimo et al., 2025; Gelens et al., 2024; Imperatori et al., 2019; Potash et al., 2025b; Roberts et al., 2025; Vinck et al., 2023).

Here, we present the current neurophenomenological research as a novel approach to studying PA. To address issue (i), we adopt Temporal Experience Tracing (TET) (Gernert et al., 2024; Jachs, 2021; Jachs et al., 2022; Lewis-Healey et al., 2024a,b; Niedernhuber et al., 2024), a recent methodology that empirically charts the dynamics of subjective experience. In TET, participants retrospectively reconstruct phenomenological time series by sketching, on two-dimensional axes, how the intensity of predefined experiential dimensions fluctuated over time. This produces richer and temporally resolved trajectories that surpass the capabilities of static Likert ratings (Figure 1B). To address issue (ii), we performed a large-scale screening of theoretically motivated EEG neural markers spanning complementary levels of neural dynamics. At the local signal level, we quantified aperiodic activity (exponent and offset), permutation entropy (PE), and Lempel–Ziv complexity (LZ), indexing broadband spectral structure, temporal unpredictability, and signal complexity. At the network level, we quantified linear functional connectivity using weighted phase-lag index (WPLI) and nonlinear information sharing using weighted symbolic mutual information (WSMI). Together, these measures were used as complementary probes of local dynamics, oscillatory coupling, and nonlinear information exchange that may distinguish PA from ordinary cognition and baseline rest (Figure 2). We recorded EEG and TET from 33 TM practitioners and their corresponding matched controls (who performed a counting task) to determine phenomenological and neurophysiological differences in PA between TM practitioners and controls. Using multivariate classification analyses, we provide a comprehensive ranking of the most discriminative neural markers separating TM from controls during the experience of PA, as well as the most relevant neural markers distinguishing meditation from baseline periods.

**Figure 1:**
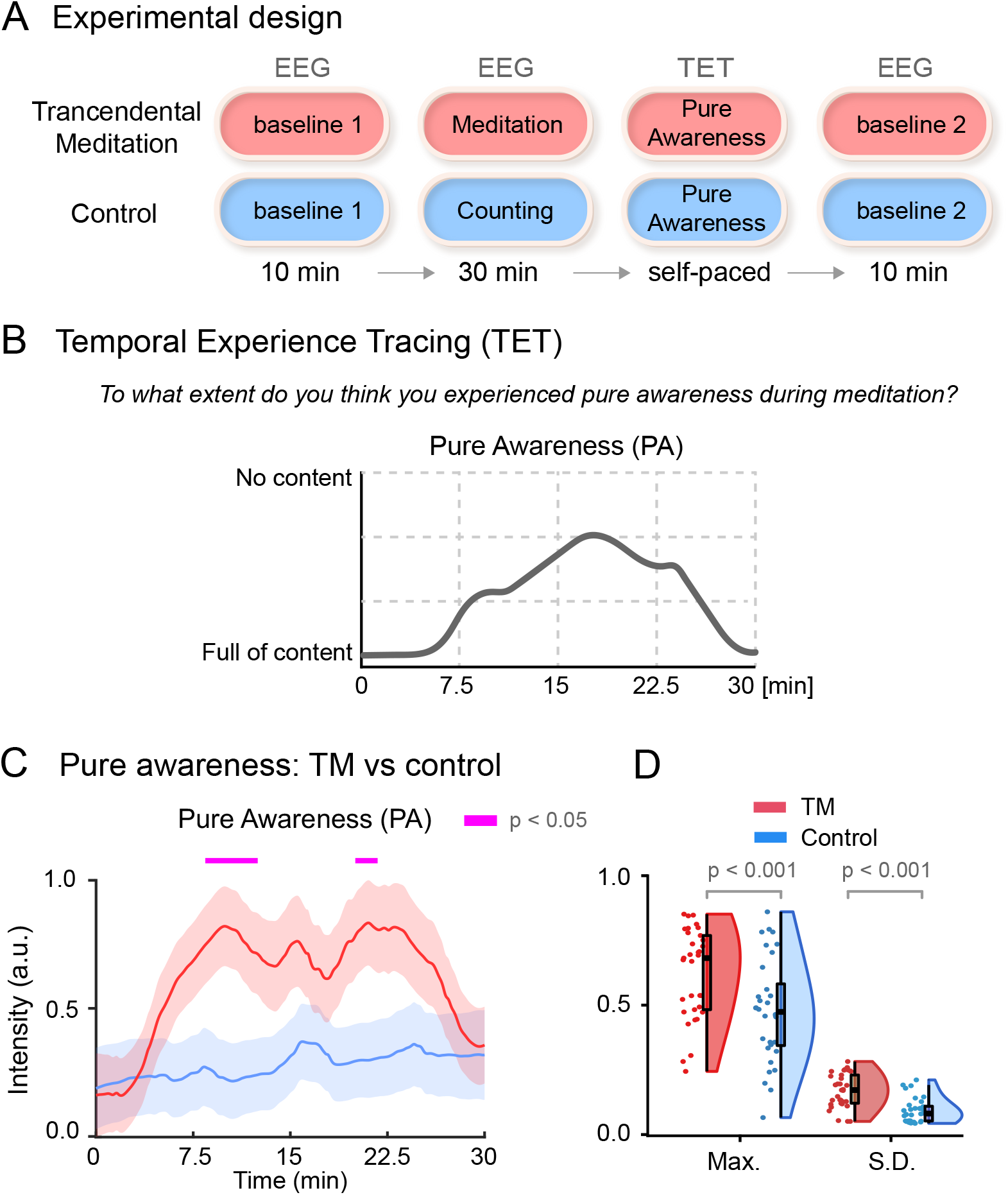
Experimental design, temporal experience tracing, and pure awareness differences. **(A)** High-density EEG from TM and control participants was recorded during a baseline condition (B1) for 10 minutes, eyes closed, without any specific instructions. Then, they practiced TM for 30 minutes (M), after being instructed to start repeating their mantra. After meditation, they were immediately instructed to report their experience of pure awareness (PA) using the TETs. Finally, a last baseline condition (B2) was recorded for another 10 mins, with no specific instructions other than keeping their eyes closed. **(B)** Pure awareness report using TET. Schematic trace representation of single-participant reporting PA during the 30-min meditation session (see Methods). **(C)** Group-level differences in PA between TM and controls. The average TET values across TM (in red) and control (in blue) participants. A cluster-based permutation test over time reveals significant temporal clusters with higher PA in TM compared to controls (see Results and Methods sections). Shaded bars indicate standard error of the mean (S.E.M.). **(D)** Maximum PA intensity and SD across time (see Results and Methods sections) were compared between the TM and control groups. Box plots display the sample median alongside the interquartile range. Individual dots represent individual PA values per subject (TM in red, controls in blue).

**Figure 2:**
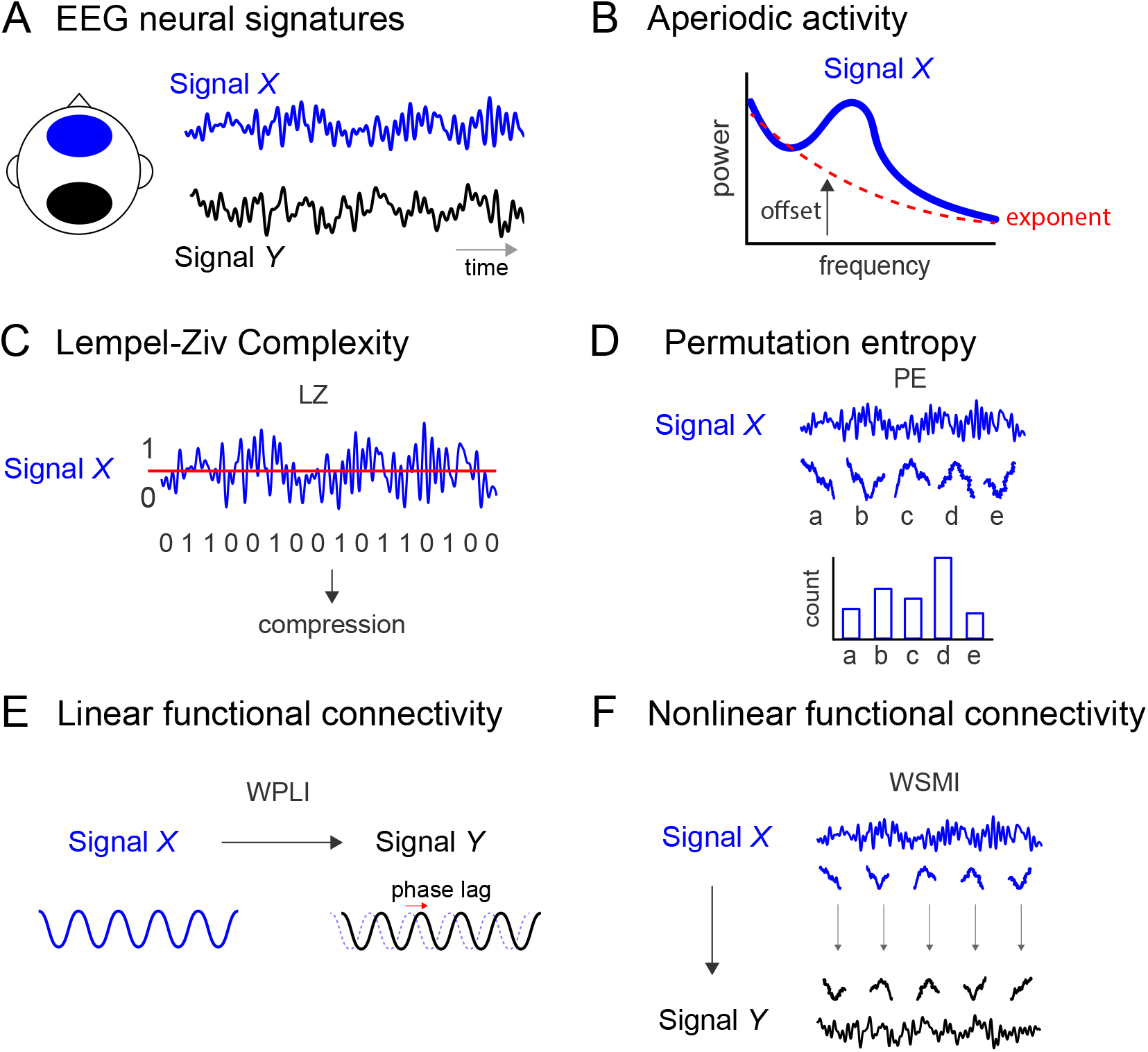
Theoretically motivated EEG neural measures. **(A)** Extracting neural features from EEG signals. In this diagram, neural signals for signals X (in blue) and Y (in black) are depicted, corresponding to two arbitrary electrodes of interest. **(B)** Aperiodic activity. **(C)** Lempel-Ziv complexity. **(D)** Permutation entropy. **(E)** Linear transformations between two correlated signals at the same frequency. The transformed signal (signal Y) is the result of a phase-lagged version of the original signal (signal X) at the same frequency (e.g., a 10 Hz signal X can result in a phase-shifted signal Y at the same 10 Hz). The WPLI (weighted phase-lagged index) captures this type of linear functional connectivity. **(F)** Nonlinear transformations cause systematic relationships across different frequencies between signal X and Y, potentially in the absence of linear transformations. Thus, non-linear connectivity measures quantify arbitrary mappings between temporal patterns in one signal (signal X) and another signal (signal Y). This type of nonlinear mapping is captured by the WSMI (weighted symbolic mutual information).

## METHODS

### Participants

Meditators and controls were male and female, aged between 18 and 50 years. Participants with no chronic mental health, sleep-related, or neurological medical problems were included in the study. In the Transcendental Meditation (TM) group (*N* = 33; 21 male; *M* = 40.3 years old, *SD* = 6.2), we selected participants with a minimum of 4 years of regular meditation practice (twice a day, 20 minutes per session; *M* = 12.9 years of practice; *SD* = 7.7). The Control group (C) included participants who had not practiced TM before (*N* = 33; 21 male; *M* = 38,9 years old; *SD* = 7.5). To control for potential age and sex-related differences between groups, we individually matched controls to meditators by sex and age (±3 years).

Sample size was determined a priori based on power considerations. Assuming a two-sided independent-samples t-test (alpha = 0.05) and a medium-to-large effect size of Cohen’s d = 0.70, consistent with group differences typically reported in EEG and neuroimaging studies comparing experienced meditators with non-meditating controls in similar paradigms (Atad et al., 2025; Vivot et al., 2020). Standard power calculations indicated that at least 28 participants per group are required to achieve 80% power. To our knowledge, our sample size is larger than most previous EEG and neuroimaging studies on TM, which typically include markedly fewer participants per group. This comparatively high N represents a key strength of the present work, increasing the precision and robustness of our estimates of TM-related neural effects.

Meditators were recruited by the David Lynch Foundation UK (DLF UK), in collaboration with the Maharishi Foundation, from across the United Kingdom. Healthy controls were recruited through the University of Cambridge online recruitment system. All participants signed an informed consent, and the Cambridge Psychology Research Ethics Committee approved the experimental protocol.

### Temporal experience tracing (TET)

TET has previously been used to study the phenomenological dynamics of different meditation styles (Jachs et al., 2022; Lewis-Healey et al., 2024b), the temporal dynamics of stress in autism (Gernert et al., 2024), chronic pain (Niedernhuber et al., 2024), and the neural and experience dynamics of medium- and high-dose DMT (Lewis-Healey et al., 2024a). With the TET methodology, the dimensions of subjective experience most relevant to the phenomenon under study are identified. These dimensions are subcomponents of the contents of consciousness relating to the affective, attentional, or motivational aspects of experience. After completing a session on the particular phenomenon under study, participants provide a retrospective trace of the intensity of each specific dimension. In the current study, traces were hand-drawn on printed sheets containing two-dimensional axes, with time on the x-axis and subjective intensity on the y-axis. The procedure was retrospective rather than online, and participants completed the traces in a self-paced manner immediately after the 30-minute meditation or counting period. Although six experiential dimensions were collected in the broader protocol, the present study focuses specifically on the pure awareness (PA) dimension. Below is the definition of pure awareness, which the participants were asked to trace as depicted in Figure 1B:

#### Pure awareness

To what extent do you think that you experienced pure awareness during meditation/counting?

- Higher level of pure awareness = an experience free from content defining everyday working experiences (e.g., internal thoughts, perception of the external world), absence of time, space, and body sensation.
- Lower level of pure awareness = normal waking consciousness with a variety of internal and external content.

### Experimental design and procedure

#### Transcendental Meditation

In the meditation session, participants began with a 10-minute baseline (baseline 1; B1), during which they were instructed to keep their eyes closed without reciting the mantra. The participants were then instructed to start mentally repeating the mantra with their eyes closed and perform 30 minutes of TM (meditation condition). After meditation (or counting), the participants were introduced to the process of completing the TET reports, accompanied by an explanation of the phenomenological dimension of pure awareness (PA). Importantly, the TET instructions were given only after the task period had ended. This was done to avoid imposing an explicit metacognitive or monitoring demand during TM itself, which could have interfered with the effortless quality of the practice. After completing the TETs, the participants performed a second baseline condition of 10 minutes (baseline 2; B2), during which they were again instructed to keep their eyes closed without repeating the mantra (Figure 1A). The experimenter indicated the beginning and end of each condition.

#### Non-meditative control task

A control task was developed to involve participants in cognitive processes different from meditation but with similar automaticity and decreased cognitive control as during mantra repetition performed by the TM group. This active control was chosen to preserve key procedural features of the TM session, including eyes-closed posture, internal task engagement, minimal motor output, and self-paced execution, while differing in a principled way from TM in its phenomenological profile. The control task was a counting task in which participants were instructed to mentally count in increments of 1 with their eyes closed, at their own pace, and to restart the count at any time if they lost track. The participants completed the control task after 30 minutes (Figure 1A). Counting was preceded and followed by a baseline condition of 10 minutes (B1 and B2, respectively) in which participants had no instruction except to keep their eyes closed and stay awake (Figure 1A). The experimenter indicated the beginning and end of each condition.

### TET data preprocessing

TET data were converted to vectors using a semi-automatic custom MATLAB script, as employed in previous studies (Gernert et al., 2024; Jachs et al., 2022; Lewis-Healey et al., 2024a,b; Niedernhuber et al., 2024). The script converted the image coordinates of the TET data to graph coordinates based on the orientation of the x- and y-axes. The phenomenological dimension of PA was manually checked to ensure the vectors were similar to the TET traces themselves. After conversion, the phenomenological dimension was concatenated column-wise, creating a 1xN TET matrix for each condition per participant, where N is the number of data points generated for that session.

### TET data analysis

Next, we performed two analyses on the generated PA time series to (i) compare the maximum PA value across groups, and (ii) compare the temporal variability of PA within participants.

#### Maximum PA value

For each participant, we identified the global maximum of their PA time course. To obtain a robust estimate of this peak, we extracted a fixed-length 1-minute window (15,000 samples) centered on the maximum. The mean PA value within this peak-centered window was then computed, yielding a peak-window average PA score per participant. Group differences were statistically assessed using Welch’s independent-samples *t*-test, chosen because it provides reliable inference under unequal group sizes and does not assume equal population variances.

#### Within time course variability

To quantify temporal variability in PA, we computed the standard deviation of the PA time series across all time points for each participant, yielding a single variability index per participant. Group comparisons of these variability indices were again carried out using Welch’s *t*-test for the same reasons outlined above.

For the group comparison of PA values, we employed a non-parametric framework to address the multiple-comparisons problem, with temporal clustering implemented in FieldTrip (Maris et al., 2007). The process required calculating t values at each time point, followed by clustering adjacent t values; this was repeated 1000 times, with recombination and randomized resampling before each repetition. This Monte Carlo method generated a non-parametric estimate of the p-value representing the statistical significance of the initially identified cluster. The cluster-level t value was calculated as the sum of the individual t values at the points within the cluster. Cluster threshold: 0.05

### EEG data preprocessing

EEG signals were recorded using a 128-channel HydroCel Geodesic Sensor Net (GSN 128 1.0) and a GES400 Electrical Geodesics amplifier, sampled at 250 Hz with NetStation software (EGI, USA). During recording and analysis, the average across electrodes was used as the reference electrode. Two bipolar derivations were designed to monitor vertical and horizontal ocular movements. Eye movement contamination (blinks were rare as eyes were closed, vertical and horizontal saccades or slow movements were also infrequent), and muscle artifacts (i.e., cardiac and neck movements) were removed from the data before further processing using an independent component analysis (ICA) (Delorme and Makeig, 2004). All conditions yielded at least 96% of artifact-free trials. Trials that contained voltage fluctuations exceeding ±200 µV and transients exceeding ±100 µV were excluded. No low-pass or high-pass filtering was performed during the pre-processing stage. The EEGLAB Matlab toolbox was used for data pre-processing and pruning (Delorme and Makeig, 2004).

### Single-electrode EEG neural measures

#### Aperiodic exponent and offset

We quantified the aperiodic component of the EEG power spectrum—often termed ‘1/f’ or ‘scale-free’ activity by parameterizing the spectrum into a broadband (aperiodic) background and narrowband (periodic) oscillatory peaks (Donoghue et al., 2020b). Following established approaches (i.e., FOOOF), the aperiodic background was modeled in log–log space as a linear function of frequency, optionally with a knee. At the same time, oscillatory peaks were fit as Gaussians and removed before estimating broadband parameters. Two parameters summarize the aperiodic background: (i) the exponent (slope), which captures how power decays with frequency (steeper, more negative slopes indicate relatively greater low-frequency power), and (ii) the offset (intercept), which reflects the overall broadband power level (a vertical shift of the spectrum) independent of narrowband oscillations. These parameters were extracted from artifact-cleaned spectra over predefined frequency ranges, excluding line-noise bands, and used as features in subsequent analyses. Previous work has shown that separating aperiodic from periodic activity improves the interpretability of spectral measures and mitigates band-ratio confounds. The exponent has been linked to age- and state-related changes, and in modeling work, to shifts in the synaptic excitation-inhibition balance, while the offset indexes broadband power differences between conditions or individuals (Ameen et al., 2024a,b; Donoghue et al., 2020a,b; Voytek et al., 2015).

#### Lempel-Ziv complexity (LZ)

LZ complexity (Lempel and Ziv, 1976; Ziv and Lempel, 1978) is a robust measure that has been used extensively in the cognitive neuroscience of consciousness (Atad et al., 2025; Casali et al., 2013; Lewis-Healey et al., 2024b; Pascovich et al., 2022; Schartner et al., 2015, 2017). In brief, the electrophysiological signal is first binarized using a median split, with values above the median coded as 1 and those below as 0 (Figure 2C). A sequential parsing algorithm then traverses the binary string to detect previously unseen substrings, thereby assembling a ‘dictionary’ of patterns. The number of unique entries serves as the raw complexity estimate, which we normalize by dividing by the complexity obtained from a randomly shuffled version of the same data. We computed LZ separately for each channel in the broadband signal (0.5-40 Hz), and the resulting values were passed to the classification analyses (see below).

#### Permutation entropy (PE)

PE (Bandt and Pompe, 2002) is a robust method for comparing electrophysiological signals between states of consciousness (Frohlich et al., 2022; King et al., 2013; Sitt et al., 2014) and distinguishing between different types of behavior (e.g., periodic, chaotic, or random). One key feature of the method is its robustness to low signal-to-noise ratios compared to other similar methods (Bandt and Pompe, 2002). The basic principle of this method is to transform the time signal into a sequence of symbols before estimating entropy (Figure 2D). The transformation is made by considering consecutive sub-vectors of the signal of size n. These sub-vectors can be made of either consecutive elements or of elements separated by τ samples (where τ is an integer). The τ parameter thus defines a broad frequency-specific window of sensitivity for this measure. Since using τ values greater than 1 induces aliasing, the signal was low-pass filtered before PE calculation to maintain the frequency-band specificity of each measure (Sitt et al., 2014). Each sub-vector of length n is associated with a unique symbol, based solely on the ordering of its n signal amplitudes. Given the parameter *n*, there are n! possible vectors, corresponding to distinct categories of signal variations. After the symbolic transform, the probability of each symbol is estimated, and PE is computed by applying Shannon’s classical formula to the probability distribution of the symbols. In our case, we computed PE for every electrode and in every trial, using parameters n = 3 and τ = [64 ms (∼1-3 Hz), 32 ms (∼4-7 Hz), 16 ms (∼8-14 Hz), and 8 ms (∼15-29 Hz)], corresponding to PE delta, PE theta, PE alpha, and PE beta, respectively. The resulting values were passed to the classification analyses (see below).

### Between-electrode EEG measures: functional connectivity

#### Weighted phase lag index (WPLI)

The WPLI (Figure 2D), a widely used measure of phase coherence in EEG data (Canales-Johnson et al., 2020b, 2021a; Imperatori et al., 2019; Olivares et al., 2025; Potash et al., 2025b; Vinck et al., 2011), measures the extent to which phase angle differences between two time series *x*(*t*) and *y*(*t*) are distributed towards positive or negative parts of the imaginary axis in the complex plane (similar to the PLI). The underlying idea is that volume-conducted activity accounts for the highest proportion of detected 0° or 180° phase differences between signals. Therefore, only phase angle distributions that predominantly fall on the positive or negative side are considered to obtain a conservative estimate for real, non-volume-conducted activity. The PLI is the absolute value of the sum of the signs of the imaginary part of the complex cross-spectral density Sxy of two real-valued signals *x*(*t*) and *y*(*t*) at a time point or trial t. While PLI is already insensitive to zero-lag interactions, the weighted Phase-Lag Index (Vinck et al., 2011) further addresses potential confounds caused by volume conduction, by scaling contributions of angle differences according to their distance from the real axis, as almost ‘almost-zero-lag’ interactions are considered as noise affecting real zero-lag interactions:

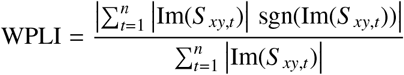

The WPLI is based solely on the imaginary component of the cross-spectrum and is thus more robust to noise than coherence, since uncorrelated noise sources increase the signal power. WPLI was computed using the Fieldtrip toolbox (multi-taper method, fast Fourier transform, single Hanning taper, 0.5 Hz frequency resolution). For each frequency range (i.e., 1-3 Hz, 4-7 Hz, 8-14 Hz, 15-29 Hz; corresponding to WPLI delta, WPLI theta, WPLI alpha, and WPLI beta, respectively) and epoch. The resulting values for each electrode pair were passed to the classification analyses.

#### Weighted symbolic mutual information (WSMI)

We quantified information sharing between electrodes using weighted symbolic mutual information (WSMI). This index estimates the extent to which two EEG signals exhibit non-random joint fluctuations (Figure 2F). WSMI has been proposed as a measure of neural information sharing in the cognitive neuroscience of consciousness literature (Canales-Johnson et al., 2020b; Imperatori et al., 2019; King et al., 2013; Olivares et al., 2025; Potash et al., 2025b; Sitt et al., 2014) and has three main advantages. First, it is a rapid and robust estimate of the signals’ entropy (i.e., the statistical uncertainty in signal patterns), as it reduces the signal’s length (i.e., its dimensionality) by looking for qualitative or ‘symbolic’ patterns of increase or decrease. Second, it efficiently detects highly nonlinear coupling between signals (i.e., non-proportional relationships between signals), as it has been shown with simulated data (Imperatori et al., 2019) and experimental EEG, electrocorticography (ECoG), and local field potentials (LFP) data (Canales-Johnson et al., 2020a,b; Imperatori et al., 2019; King et al., 2013; Olivares et al., 2025; Potash et al., 2025b; Sitt et al., 2014). Third, it rejects spurious correlations between signals that share a common source, thus prioritizing non-trivial pairs of symbols.

We calculated WSMI between each pair of electrodes, for each trial, after transforming the EEG signal into a sequence of discrete symbols defined by the ordering of κ time samples with a temporal separation between each pair (or τ). The symbolic transformation is determined by a fixed symbol size (κ = 3; where 3 samples represent a symbol) and the variable τ between samples (i.e., the temporal delay between samples), thus determining the frequency range in which WSMI is estimated (King et al., 2013). The frequency specificity *f* of WSMI is related to k and τ as follows:

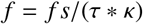

With a *f s* = 250 Hz, as per the above formula, and a *k* size of ∼3, τ = 64 ms is sensitivity to frequencies ∼1-3 Hz, τ = 32 ms (∼4-7 Hz), τ = 16 ms (∼8-14 Hz), and τ = 8 ms (∼15-29 Hz), corresponding to WSMI delta, WSMI theta, WSMI alpha, and WSMI beta, respectively. The weights were added to discard the conjunction of identical and opposite-sign symbols, which indicate spurious correlations due to volume conduction. The WSMI (in bits) can be calculated as:

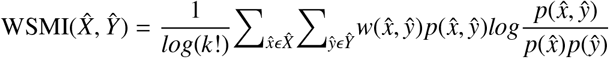

Where x and y are all symbols present in signals *X* and *Y* respectively, *w*(*x, y*) is the weight matrix, and *p*(*x, y*) is the joint probability of co-occurrence of symbol *x* in signal *X* and symbol *y* in signal *Y*. Finally, *p*(*x*) and *p*(*y*) are the probabilities of those symbols in each signal, and *K*! is the number of symbols used to normalize the mutual information (MI) by the signal’s maximal entropy. The resulting values for each electrode pair were then passed to the classification analyses.

### Implementation and evaluation of classifiers for neural markers

#### Model and Training

A multivariate Random Forest (RF) classifier was implemented within a scikit-learn pipeline consisting of feature standardization followed by classification. The feature set included spectral (aperiodic exponent and offset), functional connectivity (WPLI and WSMI), entropy (PE), and complexity (LZ) measures. The model’s hyperparameters (number of trees, maximum depth, split criterion, feature sub-sampling strategy, minimum samples per leaf, and bootstrap setting) were optimized using an exhaustive grid search. To prevent information leakage, the subject’s identity was used as the grouping variable in all cross-validation. Model training and evaluation followed a nested cross-validation scheme: the outer loop employed five-fold group cross-validation to estimate generalization, while the inner loop performed five-fold cross-validation on the training split to select hyperparameters. To reduce variance from any single partitioning, the outer loop was repeated 100 times with permuted fold assignments, and the best estimator from each repetition was retrained on the training fold and evaluated on the corresponding held-out test fold. Given 128 EEG channels, the final per-trial feature vector was high-dimensional. Specifically, single-channel measures contributed 256 aperiodic features (128 exponents + 128 offsets), 128 LZ features, and 512 PE features (128 electrodes × 4 bands). Pairwise measures were computed across 8,128 unique electrode pairs, yielding 32,512 WPLI and 32,512 WSMI features (8,128 pairs × 4 bands each), for a total of 65,920 features per trial. Because this high-dimensional setting is potentially susceptible to overfitting, all cross-validation splits were defined at the subject level; hyperparameter optimization was restricted to the inner loop; preprocessing was fit on training data only within each fold; and performance was always estimated on held-out subjects in repeated outer-loop validation.

#### Performance metrics

For each repetition, we calculated the overall accuracy, balanced accuracy (to account for class imbalance), and class-specific precision, recall, and F1-scores. Precision was defined as the proportion of true positives among predicted positives, recall as the proportion of true positives among actual positives, and F1 as the harmonic mean of precision and recall. Confusion matrices were constructed to visualize misclassification patterns. Model performance was summarized across repetitions by reporting median values and variability estimates.

#### Feature importance

To identify the neural markers contributing most to classification, RF feature importance values were extracted from the best estimator of each repetition. Importance values were analyzed both at the single-feature level and after grouping features by family (i.e., aperiodic, linear functional connectivity, nonlinear functional connectivity, entropy, and complexity) and frequency band. Grouped importance was averaged across repetitions to provide a stable estimate of feature relevance. Because many EEG-derived features are correlated across electrodes, frequency bands, and feature families, RF feature-importance values should be interpreted cautiously. In particular, impurity-based importance does not uniquely separate shared variance across correlated predictors and therefore should not be interpreted as a measure of unique mechanistic contribution. In the present study, feature importance is used primarily to summarize which feature families most consistently contributed to discrimination across repeated held-out evaluations.

#### Contrasts

Across all analyses, the same RF pipeline was applied; what differed was the subset of participants and conditions used to construct the feature matrix and labels. Three contrasts were defined. First, to test for state-expressed trait differences, we compared meditation epochs between TM practitioners and controls (Contrast 1: Meditation vs. Counting [TM vs. Controls]). Second, to isolate state effects within experienced practitioners, we compared meditation and baseline epochs from the same TM participants (Contrast 2: Meditation vs. Baselines within TM). Finally, to establish a benchmark in individuals without training, we compared counting and baseline epochs in the control group (Contrast 3: Counting vs. Baselines within Controls). For each trial, all features were computed separately at each electrode (for single-channel measures) and each electrode pair (for functional connectivity measures), so that the input to the classifier was a high-dimensional feature vector capturing the spatial pattern of neural markers across the scalp and their pairwise interactions. Thus, in the present context *multivariate* refers primarily to classification based on these spatially distributed configurations of neural signatures, rather than on single global averages per condition.

### Complementary topographical analyses of univariate neural-marker differences

To complement the multivariate classification analyses with a spatially resolved inferential test, we performed univariate topographical comparisons of the main neural markers across the principal contrasts examined in the study. We chose this approach because the RF classifier was trained on spatially distributed multichannel patterns, and therefore, the contribution of a given marker may depend on its topographical configuration rather than on a simple global increase or decrease in its mean value across electrodes. For each marker, we computed scalp topographies for the contrasts TM - B1, Control - B1, and TM - Control during the task period, as well as the double-difference contrast (TM - B1) - (Control - B1). Statistical significance was assessed using cluster-based permutation testing across electrodes, controlling for multiple comparisons while preserving the spatial structure of the scalp pattern. These analyses were intended as a complementary interpretive step and not as a substitute for the multivariate classification framework (Figure S1)

### Software and analyses

Statistical and signal processing analyses were performed using MATLAB (2023a), Python, and JASP statistical software (open-source).

## RESULTS

### Phenomenology of PA between TM and controls

All participants first completed a 10-minute eyes-closed baseline period, during which meditators were instructed not to meditate or repeat their mantra. Next, they engaged in a 30-minute task period (TM for meditators; a self-paced forward-counting-by-ones task for controls). After receiving instructions for the TETs, participants completed the TETs and then completed a second 10-minute eyes-closed baseline, with meditators again refraining from meditation (Figure 1A).

To capture the temporal dynamics of PA using the TET approach (Figure 1B), we performed a cluster-based permutation test comparing PA intensity between TM and controls (Figure 1C). We found two significant (*p* < 0.05) clusters showing higher PA for meditators compared with controls (first cluster: 8.32 min to 12.53 min, Cohen’s *d* = 0.57; second cluster: 20.00 min to 21.45 min, Cohen’s *d* = 0.53). As a complementary analysis to compare the maximum PA intensity between groups, we extracted the maximum PA value from the 30-minute TET charts for meditators (TM) and non-meditators (control) and compared the means between groups. We observed significantly higher PA in TM (*M* = 0.697, *SD* = 0.209) compared with controls (*M* = 0.523, *SD* = 0.241; *t* = 3.10, *p* < 0.001, mean difference = 0.174, 95% *CI* [0.062, 0.286], Cohen’s *d* = 0.771; Figure 1D). Lastly, to investigate the temporal variability of the PA experience over the 30-min period (Figure 1A), we calculated the standard deviation (*SD*) of the PA traces per participant in the TM and control groups. We then, in a separate contrast, statistically compared these SD values between the TM and control groups and found significantly greater temporal variation in TM (*M* = 0.154, *SD* = 0.079) compared with controls (*M* = 0.063, *SD* = 0.057; *t* = 5.36, *p* < 0.001, mean difference = 0.091, 95% *CI* [0.057, 0.126], Cohen’s *d* = 1.32; Figure 1C). These results suggest that the intensity and variability of pure awareness dynamics increase during the transcendental meditation period compared to performing mental counting in ordinary wakefulness. Thus, the results are consistent with the notion that during the meditation period, the phenomenology of TM is characterized by increased PA relative to normal wakefulness.

After having established self-reported PA differences between TM and controls, our objective was to test the second core idea of TM proponents regarding PA, that is, rapid, effort-independent access to PA, conceptualized as ‘automatic self-transcendence’ (Travis, 2001), is *independent* of years of meditation practice. This notion implies that both short- and long-term practitioners can, in principle, experience the same level of pure awareness in a single sitting. To empirically test this idea, we correlated the PA values obtained from TM practitioners with their years of practice and expected no association. Consistent with the prediction of the TM framework, we found no significant association between years of meditation practice and PA [PA intensity (maximum): Pearson’s *r* = -0.13; *p* = 0.445; PA variability (*SD*): Pearson’s *r* = -0.10; *p* = 0.550)].

### Extracting theoretically motivated EEG neural measures

To better characterize the neural dynamics during periods where PA was reported, we computed and performed multi-variate classification analyses on theoretically motivated EEG neural measures (Figure 2). At the level of single-electrode dynamics, we computed (i) the aperiodic (1/f) exponent, (ii) Lempel–Ziv complexity (LZ), and (iii) permutation entropy (PE), providing insights into local EEG neural dynamics (see Methods).

In brief, (i) parameterizing power spectra into aperiodic components (Donoghue et al., 2020b) is one of several approaches (Canales-Johnson et al., 2021b; Roberts et al., 2025) to isolate a broadband background whose exponent (slope) and offset (intercept) vary with age, arousal, and task demands. Modeling studies have linked the exponent to shifts in cortical excitation–inhibition balance, thereby enhancing the interpretability of spectral power beyond classic frequency bands (Donoghue et al., 2020b) (Figure 2B); (ii) LZ applied to EEG signals estimates the rate at which novel information patterns appear in a signal and is related to algorithmic complexity, making it sensitive to the richness of neural activity in conscious states and perceptual tasks (Canales-Johnson et al., 2023, 2020b; Lewis-Healey et al., 2024b; Pascovich et al., 2022; Schartner et al., 2015, 2017) (Figure 2C); and (iii) PE quantifies the temporal ordering of amplitude patterns and is grounded in dynamical systems theory (Bandt and Pompe, 2002), tracking neural signal unpredictability while maintaining robustness to noise (Frohlich et al., 2022; King et al., 2013; Sitt et al., 2014) (Figure 2D).

At the level of global EEG neural dynamics, functional connectivity measures can be categorized into those that predominantly capture (i) linear and (ii) nonlinear functional connectivity (Imperatori et al., 2019; Vinck et al., 2025, 2023). (i) WPLI emphasizes consistent non-zero-lag phase relationships, thereby reducing sensitivity to volume conduction and noise relative to traditional coherence, which makes it a principled marker of large-scale phase coherence in EEG (Vinck et al., 2011, 2023) (Figure 2E). In contrast, (ii) WSMI estimates shared information between symbolized EEG signals, capturing distributed, nonlinear dependencies that index global information sharing in alertness and meditation states (Canales-Johnson et al., 2020b; Imperatori et al., 2019; King et al., 2013; Potash et al., 2025b; Sitt et al., 2014) (Figure 2F). Therefore, combining linear and nonlinear functional connectivity measures provides a complementary assessment of global dynamics: WPLI for rhythmic coupling and WSMI for arrhythmic nonlinear information exchange across EEG electrodes.

### Neural dynamics of PA distinguishing meditation from control

To determine whether these theoretically motivated neural signatures contain information relevant to distinguishing meditation from counting within the TM and control groups, we performed multivariate Random Forest (RF) classification. The RF classifier trained on meditation and counting EEG trials achieved an overall precision of 65%, corresponding to a balanced accuracy of 65% (Figure 3A, Table S1, and Table S2). Meditators were correctly identified in 67% of cases, while controls were correctly classified in 62% of cases. Misclassifications were distributed so that 33% of the meditators were misclassified as controls, and 38% of the controls were misclassified as meditators.

**Figure 3:**
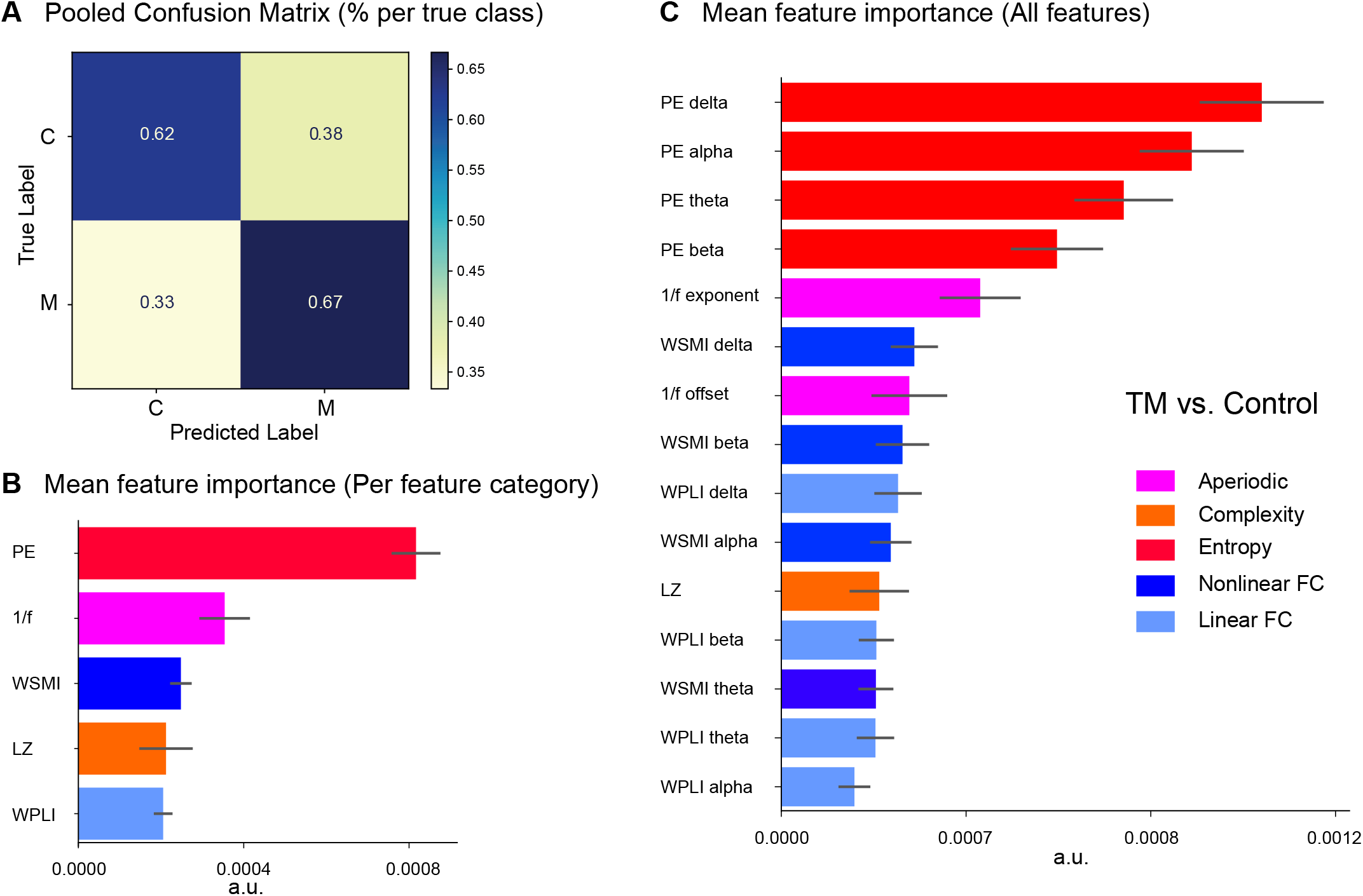
Multivariate Random Forest (RF) classification contrasting meditation (TM) and counting (control). **(A)** Pooled confusion matrix showing classification performance across 100 outer cross-validation repetitions (i.e., percent of true class samples correctly/incorrectly labeled). **(B)** Mean feature importance aggregated by feature category, averaged across repetitions. Single-electrode features: Permutation entropy (PE; in red), Aperiodic activity (1/f; in pink), Lempel-Ziv complexity (LZ; in orange). Functional connectivity features (between-electrodes): Nonlinear functional connectivity computed with weighted symbolic mutual information (WSMI; blue), and Linear functional connectivity computed with weighted phase-lag index (WPLI; light blue). **(C)** Mean feature importance (in arbitrary units; a.u.) at the level of individual features, color-coded by feature category (Entropy: PE in delta, theta, alpha and beta bands; Aperiodic activity: exponent and offset; Complexity: LZ, Linear functional connectivity: WPLI in delta, theta, alpha and beta bands; and Nonlinear Functional Connectivity: WSMI in delta, theta, alpha and beta bands). Error bars denote the standard error of the mean (S.E.M.) across repetitions. Together, the results highlight the predominance of temporal entropy measures (i.e., PE) in driving group separability, followed by aperiodic parameters and, to a lesser extent, functional connectivity and complexity features.

These results indicate that, when comparing meditation with counting, the selected EEG neural markers provided sufficient information to distinguish practitioners from controls with robust reliability, with meditators showing higher classification consistency than controls. These findings suggest that the meditation condition, in which meditators exhibited a higher phenomenology of PA compared to the control group, is associated with a more distinctive neural signature.

### Feature relevance of PA distinguishing meditation from control

To identify in more detail the contribution of each neural marker to separating meditation from counting (control), we ranked them by their relevance to classification, first at the level of feature families (Figure 3B) and then at the level of individual features (Figure 3C). Inspection of feature importance indicated that PE contributed most strongly to group classification, with the highest mean values observed in the delta, theta, and alpha frequency bands. These were followed by PE beta, which also showed above-average importance. The exponent and offset of the aperiodic spectral activity contributed intermediate values. At the same time, the nonlinear functional connectivity measure WSMI and the linear functional connectivity measure WPLI across the delta, theta, alpha, and beta bands, as well as LZ complexity, showed relatively lower importance. Overall, the temporal entropy features accounted for the majority of discriminative power, with spectral slope and nonlinear functional connectivity measures providing more modest contributions (Table S1 and Table S2).

PE neural markers consistently outperformed all other categories, highlighting the information diversity of temporal signals on multiple scales as the most reliable discriminator of group membership in this task context. This finding is consistent with previous evidence that meditation practice is associated with increased entropy and dynamical richness of neural activity, reflecting a broadened repertoire of brain microstates (Vivot et al., 2020). The secondary contribution of the spectral slope parameters (exponent and offset) suggests that alterations in the aperiodic 1/f background also carry meaningful information to distinguish meditation from counting. This result is consistent with reports linking contemplative states with shifts in the cortical excitation-inhibition balance (Colombo et al., 2019; Rodriguez-Larios et al., 2021). Taken together, these findings suggest that the most robust neural signatures of meditation during active engagement (meditation versus counting) are reflected in the temporal structure of EEG dynamics.

Importantly, these feature-importance values indicate how strongly each neural marker, as distributed across electrodes, contributes to the multivariate decision boundary between meditation and counting. They should not be interpreted as showing that PE or aperiodic neural markers are “higher” or “lower” in meditators relative to controls. Instead, the classifier exploits a spatial pattern of entropy and aperiodic dynamics across the scalp, together with weaker contributions from functional connectivity and complexity, to distinguish between groups. Importantly, the aim of this analysis was to identify which multivariate neural signatures best discriminated the contrasted states, not to infer the direction of change of each marker when considered in isolation.

To clarify the baseline-relative changes underlying the classification results, we performed complementary univariate topographical difference analyses for the main neural markers (Figure S1). These maps showed that the clearest and most spatially consistent within-condition effects were expressed in the aperiodic parameters, particularly the 1/f exponent and offset, whereas entropy and complexity measures were more heterogeneous across the scalp (Figure S1A–C). Importantly, the direct double-difference comparison between groups did not reveal robust corrected electrode-wise differences, indicating that the classifier’s separation of TM from the control condition is not driven by a few localized univariate effects, but rather by distributed multivariate patterns (Figure S1D).

### Neural dynamics distinguishing meditation from baselines

Having established the neural signatures that most clearly distinguish meditation from counting, we investigated whether these markers also distinguished meditation from non-meditation periods. This within-condition contrast is important because it anchors the meditation state to each participant’s own resting baseline, thereby reducing between-person heterogeneity and clarifying which neural signatures change relative to rest. To this end, we performed a second RF classification analysis comparing meditation with the baseline periods (pre-meditation: B1 and post-meditation: B2) in the TM group (Figure 4A, Table S1, and Table S2). The RF classifier achieved a robust separation between meditation and baselines, but struggled to distinguish between B1 and B2 (Figure 4A). Meditation epochs were correctly identified with 88% sensitivity, with only 12% misclassified as baseline. By contrast, the classification of baselines was more ambiguous. B1 trials were split between correct classification (64%) and misclassification as B2 (33%), while B2 was correctly classified in 55% of cases but mislabeled as B1 in 45%.

**Figure 4:**
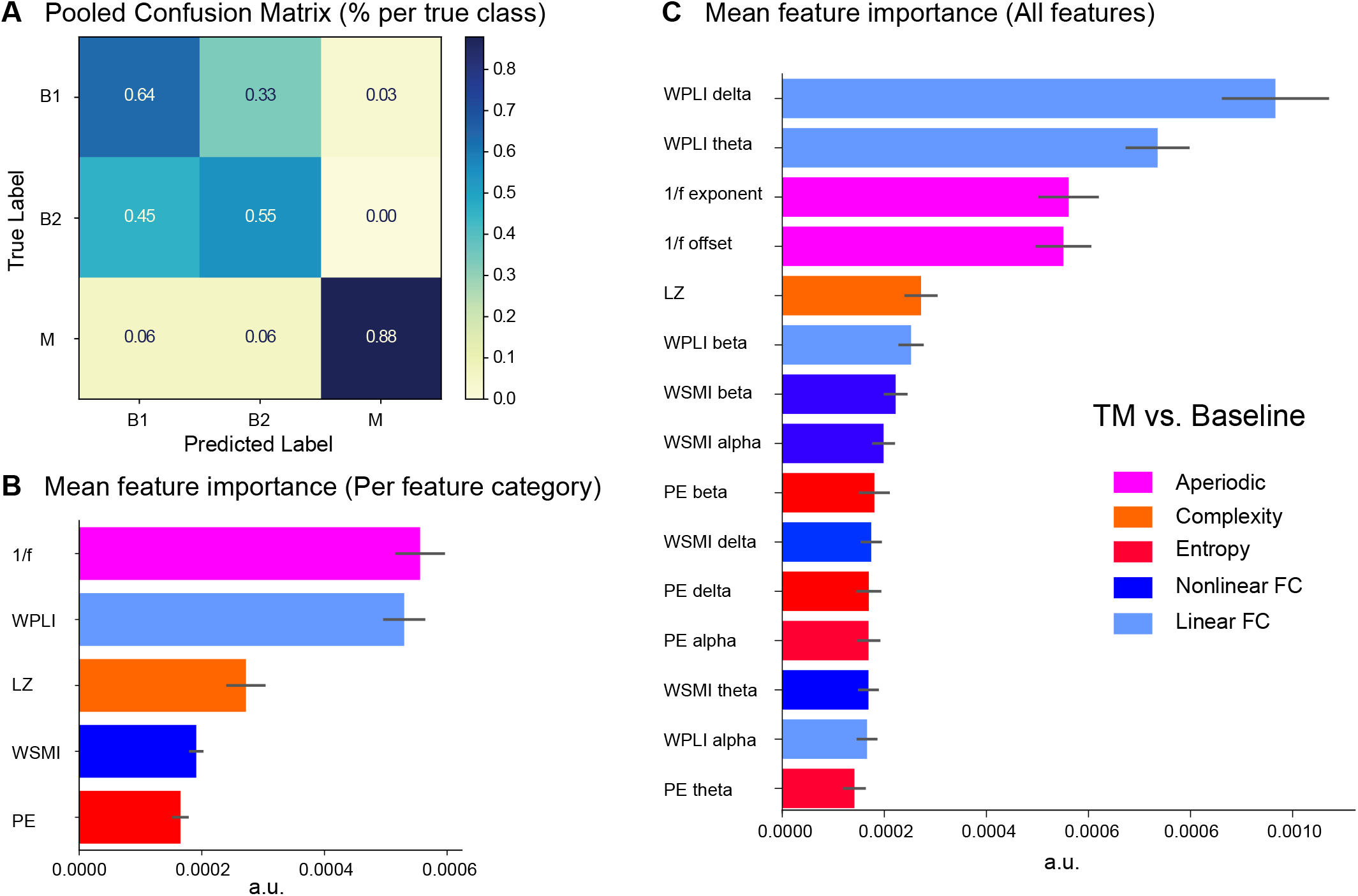
Multivariate Random Forest (RF) contrasting meditation (TM) from baselines (B1 and B2). **(A)** Pooled confusion matrix showing classification performance across 100 outer cross-validation repetitions (i.e., percent of true class samples correctly/incorrectly labeled). **(B)** Mean feature importance aggregated by feature category, averaged across repetitions. Single-electrode features: Permutation entropy (PE; in red), Aperiodic activity (1/f; in pink), Lempel-Ziv complexity (LZ; in orange). Functional connectivity features (between-electrodes): Nonlinear functional connectivity computed with weighted symbolic mutual information (WSMI; blue), and Linear functional connectivity computed with weighted phase-lag index (WPLI; light blue). **(C)** Mean feature importance (in arbitrary units; a.u.) at the level of individual features, color-coded by feature category (Entropy: PE in delta, theta, alpha and beta bands; Aperiodic activity: exponent and offset; Complexity: LZ, Linear functional connectivity: WPLI in delta, theta, alpha and beta bands; and Nonlinear Functional Connectivity: WSMI in delta, theta, alpha and beta bands). Error bars denote the standard error of the mean (S.E.M.) across repetitions. Together, the results highlight the predominance of linear functional connectivity (i.e., WPLI) and aperiodic dynamics, as well as complexity features, nonlinear functional connectivity, and entropy, to a lesser extent.

### Feature relevance distinguishing meditation from baselines

To quantify the contribution of each neural marker to classification performance between TM and baselines, we ranked the neural features by their relative importance (Figure 4B). Examination of these ranks indicated that linear functional connectivity in the low-frequency range, i.e., WPLI in the delta and theta bands, accounted for the highest discriminative power (Figure 4C). The neural markers of aperiodic activity (exponent and offset) followed with intermediate importance, with LZ complexity contributing at a comparable but slightly lower level. By contrast, the nonlinear connectivity measures WSMI and PE contributed comparatively modestly (Table S1 and Table S2). In sum, linear functional connectivity in the lower frequency bands dominated the classification analysis, aperiodic spectral parameters provided secondary relevance, and signal entropy and complexity measures added only limited incremental information. Again, the model relies on the spatial pattern of low-frequency WPLI and aperiodic parameters across electrodes and electrode pairs, rather than on global shifts in their mean values. Meditation is thus characterized by a particular topography and spatial pattern configuration of these markers relative to rest, not simply “more” functional connectivity or “less” entropy overall.

Taken together, the strong sensitivity in detecting meditation trials indicates that the meditation period engages a distinctive neural pattern that reliably diverges from both baselines. By contrast, the difficulty in separating B1 from B2 suggests the EEG of meditators at rest is highly comparable before and after meditation. These results suggest a negligible carry-over effect after TM, highlighting the specificity and transient nature of PA.

### Neural dynamics distinguishing control from baselines

After identifying the more relevant neural markers for distinguishing meditation from rest in the TM group, we evaluated how well these features separated counting from the two resting baselines in the control group (Figure 5A, Table S1, and Table S2). The RF classifier achieved a clear separation across conditions (Figure 5A). Counting epochs were correctly classified in 84% of cases, with only limited spillover into baseline labels (12% in B1; 3% in B2). Similarly, the baseline periods were also classified well: B1 was correctly classified in 84% of trials (16% mislabeled as B2), and B2 was correctly classified in 81% of trials (19% mislabeled as B1). Notably, no baseline segments were mislabeled.

**Figure 5:**
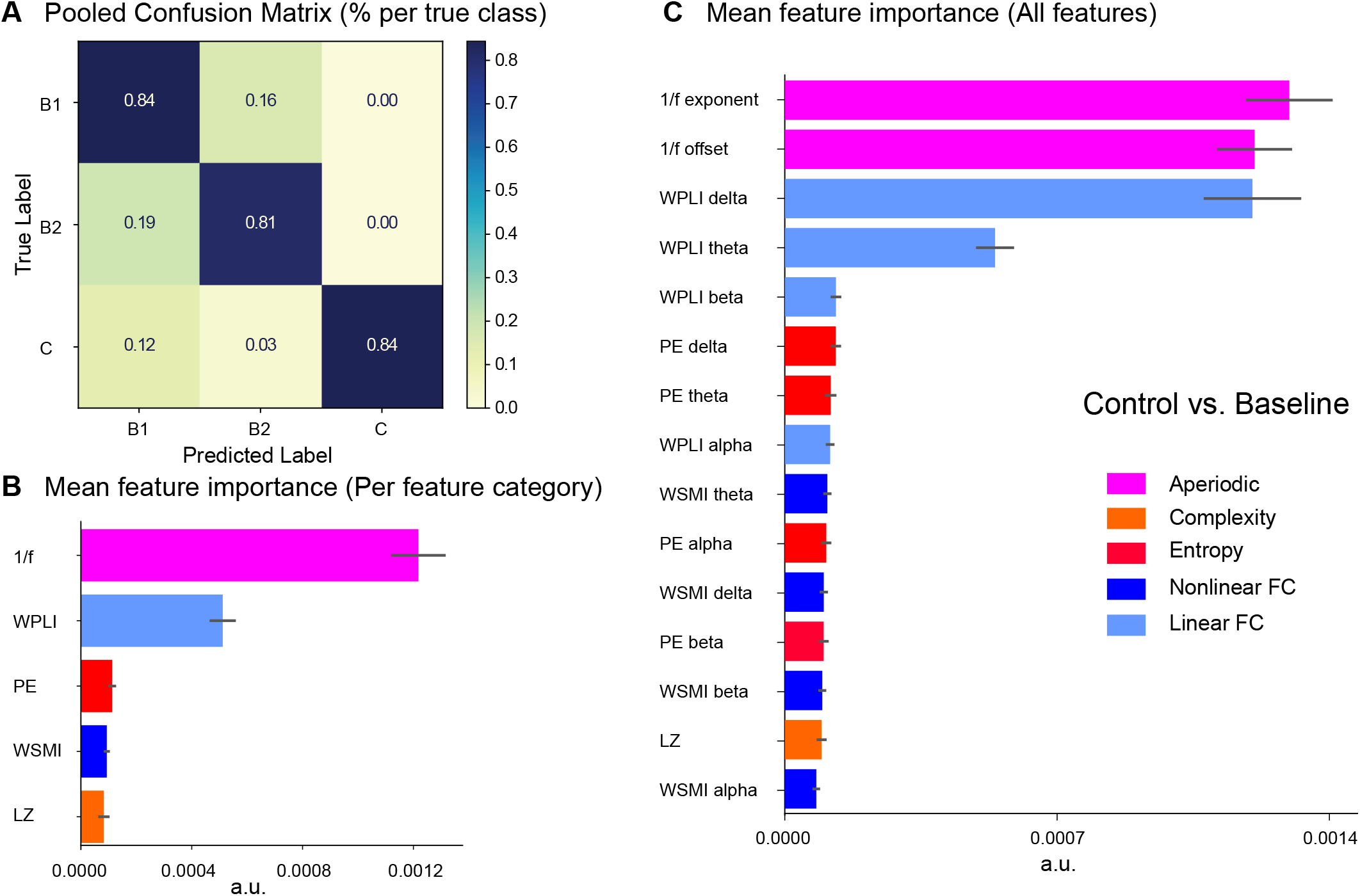
Multivariate Random Forest (RF) contrasting control (counting) from baselines (B1 and B2). **(A)** Pooled confusion matrix showing classification performance across 100 outer cross-validation repetitions (i.e., percent of true class samples correctly/incorrectly labeled). **(B)** Mean feature importance aggregated by feature category, averaged across repetitions. Single-electrode features: Permutation entropy (PE; in red), Aperiodic activity (1/f; in pink), Lempel-Ziv complexity (LZ; in orange). Functional connectivity features (between-electrodes): Nonlinear functional connectivity computed with weighted symbolic mutual information (WSMI; blue), and Linear functional connectivity computed with weighted phase-lag index (WPLI; light blue). **(C)** Mean feature importance (in arbitrary units; a.u.) at the level of individual features, color-coded by feature category (Entropy: PE in delta, theta, alpha and beta bands; Aperiodic activity: exponent and offset; Complexity: LZ, Linear functional connectivity: WPLI in delta, theta, alpha and beta bands; and Nonlinear Functional Connectivity: WSMI in delta, theta, alpha and beta bands). Error bars denote the standard error of the mean (S.E.M.) across repetitions. Together, the results highlight the predominance of aperiodic dynamics followed by linear functional connectivity (i.e., WPLI) and, to a lesser extent, nonlinear functional connectivity and entropy/complexity features.

### Feature relevance distinguishing controls from baselines

As in the previous sections, we ranked features by classification relevance to quantify their relative contributions (Figure 5B,C). The aperiodic spectral exponent and offset emerged as the strongest contributors, followed by linear functional connectivity in the lowest frequency bands (WPLI delta and WPLI theta). PE in the delta and theta ranges, as well as the remaining connectivity features, contributed comparatively less (Tables S1 and S2). As in the previous analyses, this EEG signature is expressed as a spatial configuration of spectral and connectivity features across electrodes and electrode pairs. Thus, the classifier’s performance reflects sensitivity to these multivariate spatial patterns, rather than to univariate increases or decreases in any single marker when averaged over the scalp.

Taken together, these findings demonstrate that the counting task elicits a robust EEG signature distinct from rest, primarily characterized by broadband spectral parameters and low-frequency linear functional connectivity. Crucially, the ability to differentiate between B1 and B2 indicates detectable carry-over effects from task engagement to subsequent rest, suggesting that resting activity is not merely a return to baseline levels before task engagement in the control (counting) group.

## DISCUSSION

This study used a large, rigorously controlled sample comprising 33 experienced TM practitioners and carefully age- and sex-matched controls. To our knowledge, this is the most extensive and systematically controlled neurophenomenological study of PA to date, allowing us to distinguish neural signatures of PA from potential demographic confounds. Methodologically, we advanced the field by employing Temporal Experience Tracing (TET) to study PA. We demonstrated that TM practitioners reliably report greater intensity and variability of PA, independent of years of practice, supporting the principle of automatic self-transcendence. Our large-scale neural screening and multivariate classification analyses revealed a conceptual double dissociation: relative to ordinary cognition in controls, PA is best characterized by increased entropy and scale-free neural dynamics, whereas relative to the meditators’ own baselines, PA is marked instead by enhanced low-frequency functional connectivity. In other words, PA appears as a state that is both broader and more variable than ordinary cognition, yet also more stabilized and coherent than resting baselines. Importantly, TM showed no carry-over effects, unlike controls whose task induced residual changes, highlighting the specificity and transience of PA.

Crucially, these findings do not rely on simple univariate comparisons of the average magnitude of each neural marker between conditions (e.g., meditation vs. counting). Our RF multivariate analyses operate on neural features that encode each measure at each electrode and at each electrode pair, thereby exploiting spatially distributed patterns of entropy/complexity, aperiodic dynamics, and functional connectivity. A given marker can therefore be highly informative because of the pattern it forms during PA, even if its scalp-averaged value shows little or no group difference. This emphasizes that the neural signature of PA is best understood as a multivariate spatial pattern of neural dynamics, not as “more” or “less” of any single EEG measure.

The more modest between-group classification performance relative to the within-condition baseline contrasts is not unexpected. Between-group discrimination must separate condition-related neural structure from substantial inter-individual variability, including trait-level differences in neural dynamics, idiosyncratic baseline states, and variability in task execution. By contrast, within-condition contrasts are anchored to each group’s own baseline and therefore reduce between-person heterogeneity, making state-related changes easier to detect. The weaker TM versus control classification should therefore not be interpreted as evidence of weak state effects, but rather as indicating that the discriminative signature of meditation relative to counting is distributed and embedded within broader inter-individual variability.

The strong control (counting) versus baseline classification likewise supports using counting as an active control condition. The goal of counting was not to mimic the phenomenology of TM, but to provide an internally generated, eyes-closed cognitive state with sustained engagement, self-pacing, and low overt motor demands. Its robust separation from the resting condition confirms that it is a meaningful active comparison condition. Importantly, the absence of robust univariate differences at the single-electrode level in the direct comparison between tasks suggests that the TM versus control distinction is not driven by a few large localized univariate effects, but rather by distributed multivariate patterns.

The choice of counting as an active control should also be situated relative to the broader meditation literature. Compared with passive rest, counting better matches TM in posture, eye closure, internal task engagement, low motor demands, and self-paced execution, while remaining phenomenologically distinct from the reported experience of PA. This is important because many meditation neuroimaging studies have relied primarily on passive rest as the comparison condition (Froeliger et al., 2012). Other work has used mind wandering contrasts to separate meditative attention from spontaneous internal mentation (Hasenkamp et al., 2012). Recent advanced-meditation studies have also begun to use more structured internal controls, including counting and memory-related comparison states, to avoid the interpretive limitations of rest alone (Potash et al., 2025a). Future work should therefore compare TM against multiple active and passive controls to test the specificity of the present dissociation.

### Towards a neurophenomenology of minimal phenomenal experiences

Our findings can be situated within the broader theoretical discourse on minimal phenomenal experience (MPE) as articulated by Metzinger (2020). According to this framework, MPEs represent consciousness stripped of its ordinary structural scaffolds (i.e., selfhood, temporality, spatiality) while retaining tonic alertness. PA, as reported by practitioners of TM, closely exemplifies this condition: an episode of wakefulness largely devoid of representational content, intentionally cultivated through an effortless, standardized practice.

The neurophenomenological signatures of PA uncovered in this study were characterized by increased temporal entropy and aperiodic dynamics relative to ordinary cognition, as well as stable slow-frequency phase coherence relative to rest, mapping well onto the hypothesized architecture of MPE. Modulations in the temporal entropy of the EEG signals suggest an expanded, content-reduced repertoire of neural microstates, consistent with a form of alert presence that lacks specific intentional objects. At the same time, changes in low-frequency phase coherence provide a potential substrate for maintaining a wakeful but content-reduced mode of awareness. Notably, the absence of carry-over effects in TM highlights the transience and state-specificity of PA, distinguishing it from potential trait-like effects, such as drowsiness or boredom, that might have been elicited by 30 minutes of counting in the controls. However, the present findings should not be taken to imply that minimal phenomenal content is intrinsically or uniquely associated with modulations in neural complexity or temporal entropy. Rather, our results should be contextualized within the experience of PA during TM practice and not as a universal neural definition of pure awareness or minimal phenomenal experience, or other endogenously generated absorptive states induced by sustained and volitional mental processes (Lieberman and Sacchet, 2026)

Furthermore, our findings align with an expanding literature demonstrating that jointly acquiring phenomenological reports and neural recordings enhances the characterization of the neural dynamics underlying complex conscious states (Ganesan et al., 2024; Lewis-Healey et al., 2024b; Lutz, 2002; Olivares et al., 2015; Potash et al., 2025b; Timmermann et al., 2023; Varela, 1996). While conventional Likert scales are valuable for coarsely characterizing subjective experience (Lanfranco et al., 2021), they are inherently static, temporally sparse, prone to ceiling or floor effects, and subject to recall compression, limitations that can obscure the fine-grained structure of experience. In contrast, the TET approach used here yielded subject-specific, high-resolution trajectories that captured temporal fluctuations within each subject during the experience of pure awareness.

It is worth noticing that while TET provided fine-grained phenomenological trajectories, it necessarily relied on retrospective reporting. In the case of TM, this was a methodological requirement: introducing the dimensions of the TET beforehand would have imposed a cognitive or metacognitive load, directly undermining the effortless quality essential for TM and generating pure awareness. Finally, it must be considered that control participants, unlike TM practitioners, may lack familiarity with the experiential profile of pure awareness, which could potentially introduce demand-characteristic effects in their reports. Accordingly, future work on the neurophenomenology of minimal phenomenal experiences should prioritize comprehensive, time-resolved phenomenological methods, such as TET, to leverage neural classifiers more effectively and to recover the richness and temporal organization of subjective experience that static scales systematically miss. Looking forward, future studies could broaden the scope of TET by incorporating additional phenomenological dimensions of PA, such as degrees of bodily awareness, affective tone, or selfhood, thereby enriching the neurophenomenological profile of this state. Moreover, comparing TM with other contemplative traditions that explicitly report PA, such as Dzogchen or Mahamudra practices, would enable us to test the generalizability of our findings across cultural contexts.

### Functional connectivity results and previous TM literature

Our results revealed that nonlinear functional connectivity (WSMI) provided greater discriminative power than linear phase coherence measure (WPLI) when separating TM from controls. This suggests that PA engages complex, arrhythmic information sharing across brain regions, rather than a predominance of stable oscillatory coupling. Such nonlinear dynamics capture non-proportional dependencies between signals that go beyond simple phase-locking, aligning with prior evidence that nonlinear information exchange is crucial for differentiating conscious states and contents (Canales-Johnson et al., 2020b; Imperatori et al., 2019; King et al., 2013; Potash et al., 2025b; Sitt et al., 2014; Vinck et al., 2023). Interestingly, this contrasts with earlier studies that reported increased alpha coherence as a hallmark of TM (Travis, 2001; Travis et al., 2010). Our multivariate screening, in contrast, places alpha-phase coherence among the least informative features. One likely reason is methodological: traditional coherence is highly sensitive to volume conduction, whereas WPLI explicitly controls for spurious zero-lag correlations (Vinck et al., 2011). A second issue arises in interpreting alpha coherence as a signature of inter-areal communication or integration, as recent evidence has shown that coherence is a consequence rather than the cause of inter-areal communication (Schneider et al., 2021; Vinck et al., 2025, 2023). However, phase coherence becomes more relevant when distinguishing TM from baseline, where low-frequency WPLI supports state stabilization, although primarily in the delta and theta bands. Together, these observations suggest that nonlinear functional connectivity may capture the transient aspects of PA relative to ordinary cognition. In contrast, linear connectivity reflects the stability of PA relative to its resting state.

This reinterpretation offers a significant conceptual advancement for the neuroscience of TM. Previous studies have reported increased alpha coherence as a defining neural signature of TM, linking it to heightened integration and restful alertness. Our results build on this foundation but reframe it within a modern information-theoretic perspective. By contrasting linear and nonlinear functional connectivity measures, we demonstrate that the neural signatures of pure awareness are not limited to oscillatory phase-locking in a single frequency band, but also encompass broader shifts toward nonlinear connectivity and single-electrode entropy. In this view, alpha coherence may capture a surface-level manifestation of PA, while the underlying mechanism involves deeper, more complex forms of information sharing that extend beyond rhythmic synchronization. This synthesis reconciles the classical TM literature with contemporary approaches, positioning information-theoretic measures as a more principled and sensitive index of the state of pure awareness.

A further methodological point concerns the interpretation of RF feature-importance values. Although repeated nested cross-validation reduces the likelihood that the observed rankings are driven by a single data partition, these values remain model-dependent and can be influenced by collinearity and shared information among predictors. Accordingly, the present feature rankings should be understood as indicating which neural signatures were most useful to the classifier within this dataset, rather than as uniquely identifying the mechanistic primacy of any single EEG measure.

Finally, although the study combines temporally resolved phenomenological reports with a large set of neural markers, we did not identify robust direct associations between the PA measures and the EEG features. We do not take this to imply that such a relationship is absent. Rather, it may indicate that the relevant mapping is subtle and distributed, with subjective PA depending on multivariate neural configurations that are not well captured by simple one-to-one associations between individual EEG markers and summary phenomenological variables. Establishing that mapping will require future work using multivariate and neurophenomenological modeling.

### Conclusion

Overall, this study provides the most comprehensive neurophenomenological characterization to date of PA, a paradigmatic case of minimal phenomenal experience induced through TM. By combining a large, rigorously matched sample with temporally resolved phenomenological reports and a systematic screening of theoretically motivated neural markers, we were able to distill the distinctive neural architecture of PA. Finally, it significantly expands the neurophysiological characterization of TM, which can inform future studies on well-being and the clinical population.

## Data Availability

The data and software code supporting this study’s findings are available from the corresponding author upon reasonable request, in accordance with the Department of Psychology, University of Cambridge’s data sharing policies.

## Authorship contribution

Conceptualization and methodology: A.C-J. Investigation and data collection: A.Ch-J., S.V.M., C.P., and A.C-J. Data analysis: M.E., M.D., S.V.M, A.Ch-J., and A.C-J. Visualization: M.E., M.D., S.V.M., and A.C-J. Writing and editing: A.Ch-J, S.V.M, M.E., M.D., C.P., and A.C-J. Supervision: A.C-J. Funding acquisition: A.C-J..

## Acknowledgements

We thank Dr. Evan Lewis-Healey for his contribution during the piloting of this study; Dr. Deirdre Parsons, Prof. Sanford Nidich, Dr. Mohamed Ameen, and Swami Sarvapriyananda for helpful philosophical, theoretical, and technical discussion. This study was funded by The David Lynch Foundation UK (DLF UK) through private donors. A.C-J. is funded by a Swedish Research Council Project grant (VR; 2025-03245), an ANID/FONDECYT Regular (1240899) and ANID/FONDECYT Regular (1251273) research grants.

---

**Table S1:** Descriptive statistics of neural feature categories. The table reports feature ranks, mean (M), and standard deviation (SD) of single-electrode features 1/f, PE, LZ; and between-electrode features WPLI and WSMI (functional connectivity) for the classification analysis of TM vs. counting (see Figure 3B); the classification of TM vs. B1 vs. B2 analysis (see Figure 4B); and the classification of Counting vs. B1 vs. B2 analysis (see Figure 5B).

| Feature (count) | Meditation: TM vs. Counting |  | Meditators: B1 vs. TM vs. B2 |  | Control: B1 vs. Counting vs. B2 |  |
| --- | --- | --- | --- | --- | --- | --- |
| | Rank | $M (\pm SD)$ | Rank | $M (\pm SD)$ | Rank | $M (\pm SD)$ |
| 1/f (1024) | 2 | 3.352e-04 (4.455e-04) | 1 | 5.828e-04 (4.693e-04) | 1 | 1.254e-03 (1.141e-03) |
| WPLI (1024) | 4 | 1.474e-04 (2.442e-04) | 2 | 5.734e-04 (9.649e-04) | 2 | 4.981e-04 (1.136e-03) |
| PE (256) | 1 | 1.090e-03 (1.201e-03) | 5 | 1.391e-04 (1.451e-04) | 3 | 1.136e-04 (1.148e-04) |
| WSMI (512) | 3 | 1.853e-04 (3.046e-04) | 4 | 1.652e-04 (1.579e-04) | 4 | 9.640e-05 (1.105e-04) |
| LZ (128) | 5 | 1.382e-04 (2.347e-04) | 3 | 1.816e-04 (1.494e-04) | 5 | 9.432e-05 (1.073e-04) |

**Table S2:**
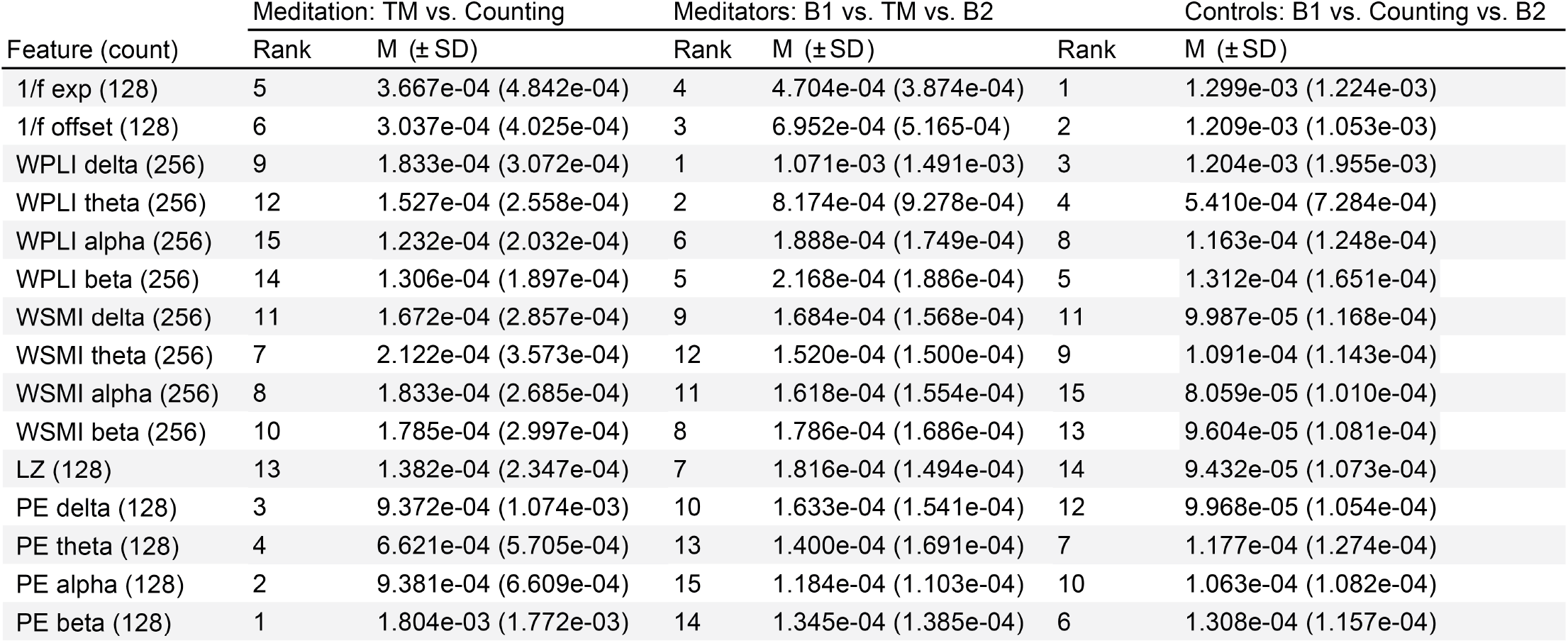
Descriptive statistics of neural feature sub-categories. The table reports feature ranks, mean (M), and standard deviation (SD) of single-electrode features 1/f exponent, 1/f offset, PE (delta, theta, alpha, and beta bands), LZ; and between-electrode features WPLI and WSMI (delta, theta, alpha, and beta bands) for the classification analysis of TM vs. counting (see Figure 3C); the classification of TM vs. B1 vs. B2 analysis (see Figure 4C); and the classification of Counting vs. B1 vs. B2 analysis (see Figure 5C).

**Figure S1:**
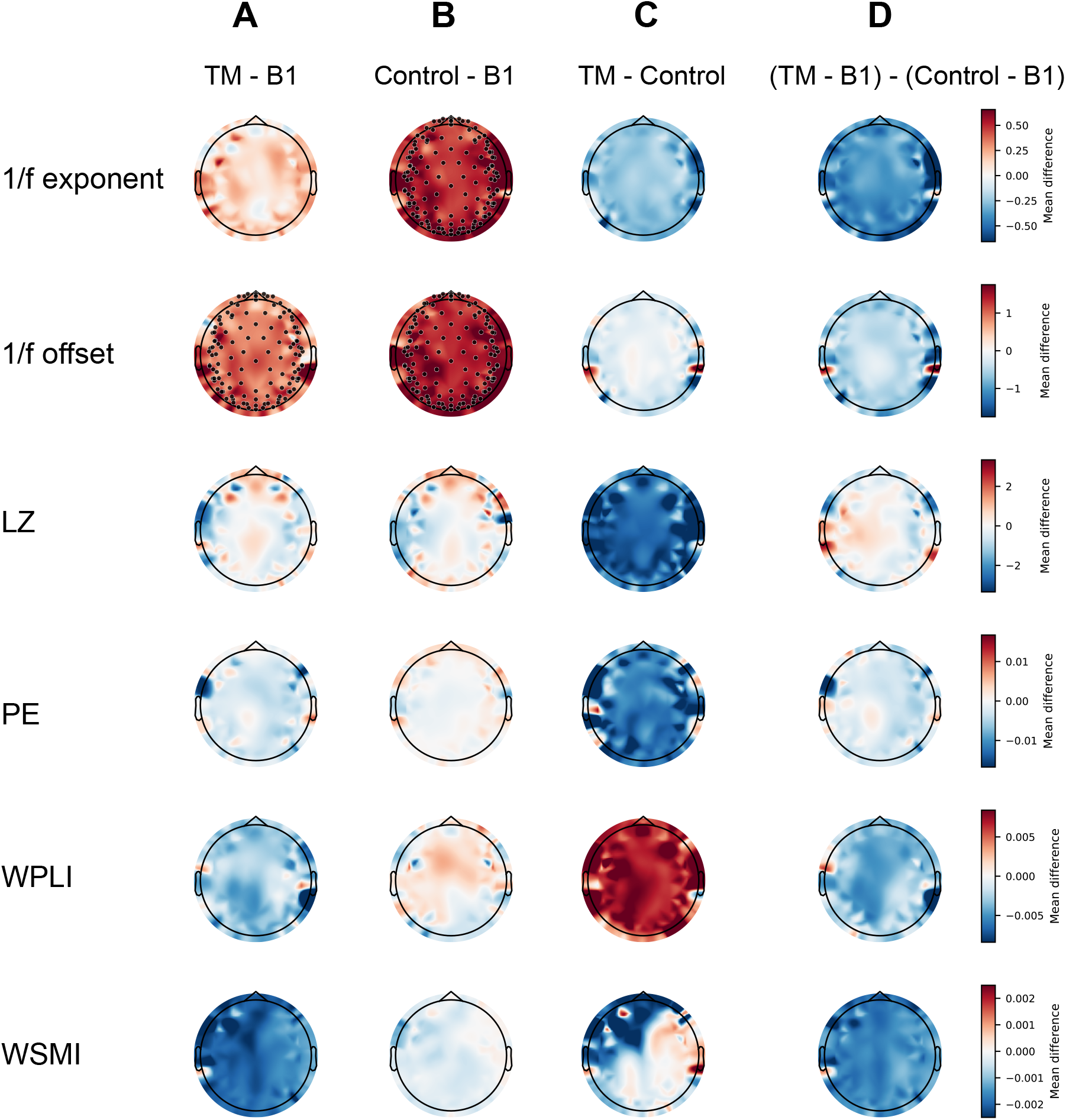
Topographical maps of univariate neural marker differences. Rows show the six neural markers studied: 1/f exponent, 1/f offset, LZ, PE, WPLI, and WSMI. Columns show A, TM B1; B, Control B1; C, TM Control during the task period; and D, (TMB1)(ControlB1). Colors represent mean differences at each electrode. Black markers denote electrodes that contributed to significant corrected effects after a cluster-based permutation test (p < 0.05). The clearest baseline-relative changes were observed for the aperiodic parameters, whereas other measures were more spatially heterogeneous. The absence of robust corrected effects in the double-difference contrast supports the view that the TM versus control classifier is driven primarily by distributed multivariate structure rather than localized univariate effects.

